# Phylogenetic analysis of *Mycobacterium bovis* Reveals Evidence Of Animal And Zoonotic Tuberculosis Transmission Between Morocco And European Countries

**DOI:** 10.1101/2024.02.09.579592

**Authors:** Hind Yahyaoui Azami, Claudia Perea, Tod Stuber, Mohammed Bouslikhane, Jaouad Berrada, Hamid Aboukhassib, Alberto Oscar Allepuz Palau, Ana C. Reis, Mónica V. Cunha, Tyler C Thacker, Suelee Robbe-Austerman, Liliana C. M. Salvador, Frederick Quinn

**Affiliations:** Department of Infectious Diseases, University of Georgia, 501 D. W. Brooks Drive, Athens, GA 30602, USA; Diagnostic Bacteriology and Pathology Laboratory, National Veterinary Services Laboratories, USDA, Ames, Iowa; Institut Agronomique et Veterinaire Hassan II, Rabat, X4GM+H88, Moroco; Faculté chouaib doukkali, Avenue Jabran Khalil Jabran B.P 299-24000, El Jadida, Morocco; Departament de Sanitat i Anatomia Animals, Facultat de Veterinària, UAB, 08193 Bellaterra, Barcelona, Spain; cE3c-Center for Ecology, Evolution and Environmental Changes & CHANGE – Global Change and Sustainability Institute, Faculdade de Ciências da Universidade de Lisboa, Campo Grande, 1749-016, Lisboa, Portugal; BioISI-Biosystems and Integrative Sciences Institute, Faculdade de Ciências da Universidade de Lisboa, Campo Grande, 1749-016, Lisboa, Portugal; School of Animal and Comparative Biomedical Sciences, The University of Arizona, Tucson, Arizona, 85721

## Abstract

Livestock production is a fundamental pillar of the Moroccan economy. Infectious diseases of cattle and other species represent a significant threat to the livestock industry, animal health, and food safety and security. Bovine tuberculosis (bTB), mainly caused by *Mycobacterium bovis* (*M. bovis*), generates considerable direct and indirect economic losses, in addition to the unknown human health burden caused by zoonotic transmission. Previous studies have suggested likely *M. bovis* transmission links between Morocco and Southern Europe, however, limitations inherent with the methods used prevented more definitive conclusions from being drawn. In this study, we employed whole genome sequencing analysis of a large set of *M. bovis* isolates to better define the phylogenetic links between strains from Morocco and neighboring countries. A total of 780 *M. bovis* sequences representing 36 countries were included in the study. The results of SNP analysis showed a close genetic relationship between *M. bovis* from Morocco and each of Spain, France, Portugal and Germany, this is supported by animal trade between Morocco and these countries, in addition to the important human migration from Morocco to Europe and North America.

Regarding zoonotic tuberculosis (TB) transmission, we were able to find genetic links between *M. bovis* isolates from cattle in Morocco and humans in Italy, Germany, and the UK. These results support our hypothesis of significant transmission of *M. bovis* from cattle to humans, which calls for further investigations of zoonotic TB transmission in Morocco and in other countries.

The fact that no *M. bovis* sequences from North Africa in the present database were classified as AF1 or AF2 clonal complexes suggests that the Sahara might play a role in preventing *M. bovis* transmission between North Africa and Sub-Saharan Africa. Our study benefits from a large sample size and a rich dataset that includes sequences from cattle, wildlife, and humans from Morocco and neighboring countries, enabling the delineation of *M. bovis* transmission routes within the animal-human interface.

## Introduction

Bovine tuberculosis (bTB) caused primarily by *Mycobacterium bovis (M. bovis)* is an important source of decreased animal health and economic distress in many countries of the world. *M. bovis* can infect a wide range of hosts, including domestic livestock, wildlife, and humans; consequently, *M. bovis* may be an important and underappreciated human health burden in countries with infected livestock and wildlife reservoirs [1]. The human health burden of *M. bovis* is poorly investigated in developing countries where bTB is endemic. Cattle-to-human transmission of *M. bovis* is primarily caused by consumption of unpasteurized milk and milk products, and secondly by inhalation of aerosols through contact with infected cattle in the farm and contact with animal carcasses and infected organs at slaughterhouses [2].

The cost of bTB to cattle farmers in Morocco has not been assessed; however, in similar settings like Ethiopia, the cost of bTB ranged from 75.2 million USD to 358 million USD per year in urban areas (where the prevalence of bTB is higher), and from 500,000 to 4.9 million USD in rural areas (where bTB prevalence is lower) [3]. It is important to note that these analyses did not consider the cost to human health.

While animal TB has been eliminated from many developed countries (e.g. Australia, Switzerland) [4,5], it remains an economic and health burden for several countries where elimination efforts are hampered by spillovers from wildlife reservoirs into livestock herds and humans (e.g. USA, UK, Canada, New Zealand) [6]. In addition, bTB is endemic in developing countries where wildlife and livestock herds are both heavily infected and in close contact (e.g. Kenya, South Africa) [7]. Although livestock herds in developed countries are routinely screened for bTB, even in developed countries, it is time- and cost-intensive to monitor wildlife reservoirs. In addition, for active surveillance, alternative diagnostic assays must be considered depending on the wildlife species. In Morocco, *M. bovis* has been isolated from the Eurasian wild boar [8]; however, investigations about its role in *M. bovis* epidemiology in that country has yet to be initiated.

Herd-to-herd transmission of *M. bovis* is variable depending on the geographic location. In countries with low bTB prevalence and strong control efforts in place (e.g., frequent bTB testing, elimination of infected animals from the herds, abattoir surveillance, and monitoring/restriction of cattle movement), transmission between herds may be relatively low (e.g. 0.001% in the USA) [9]. In developing countries, where livestock is traded without restrictions, *M. bovis* transmission between herds is common. For example, in a sero-prevalence study performed in 2012 in Morocco, bTB herd prevalence was 56% [10]. Despite strict test-and-slaughter programs, cattle movement can drive bTB dissemination between herds [9]; therefore, a combination of strategies is key in controlling prevalence of this disease.

Moreover, other animal-to-animal transmission pathways have been suggested and reported, such as vertical and pseudo-vertical transmission, in addition to auto contamination [11,12]. Environmental conditions also has been shown to impact the likelihood of animal-to-animal transmission of *M. bovis* within herds [13]. In an endemic setting like Morocco, the environment plays an important role in the transmission and persistence of *M. bovis* within and between farms. A previous study revealed an increased risk of bTB associated with intensive and semi-intensive livestock production systems, which, in Morocco, are associated with less air and sun access, increased humidity, and high animal density [10].

Phylogenetics of *M. bovis* associated with bTB cases have been increasingly investigated in the last 10 years [9,14]. Regions of deletion (RD) polymerase chain reaction (PCR), spoligotyping, and Mycobacterial Interspersed Repetitive Units-Variable Number of Tandem Repeats (MIRU-VNTR), in addition to other genotyping techniques, have been shown to be useful for *M. bovis* genetic analyses. When two or more techniques are used in conjunction, a higher genetic discrimination level between different *M. bovis* strains is achieved. Moreover, with the recent increased use of whole genome sequencing (WGS) and single nucleotide polymorphism (SNP) analysis, a broader and more in-depth knowledge about bTB transmission dynamics within and between different countries and regions is being developed.

In 2015, a molecular characterization of Moroccan *M. bovis* isolates from two slaughterhouses was performed, in which both deletion PCR (using genomic regions of difference 4 (RD4) and 9 (RD9)) and spoligotyping were used. From this study, SB0120, SB0121, and SB0265 were the most frequent spoligotypes found [15]. These have been previously reported in North African [16], European, and South and North American countries [17]. The results were supported by cattle trade data between those countries and Morocco, and the potential illegal importation of cattle from neighboring countries. Interestingly, the same study showed no spoligotypes matching to West, East, and Central Africa. The authors suggested that the desert acts like a barrier against the circulation of bTB between Sub Saharan African countries and Morocco [15]. However, deletion PCR and spoligotyping each target a single or very few genetic loci, covering less than 0.1% of the genome [18].

Whole genome sequencing is routinely used in human studies [19]and during bTB outbreaks in low prevalence settings to identify transmission sources [20,21]. Although WGS is now used sporadically in some bTB endemic settings such as Mexico [22] and Eritrea [14], and has been a valuable tool during elimination control strategies to limit potential reintroduction of *M. bovis*, it is not routinely used in most developing countries where bTB is endemic.

Based on genomic deletions and spoligotype, *Mycobacterium bovis* is classified into several groups of genotypes called clonal complexes (CC). African1 (AF1) [23], African 2 (AF2) [24], European1 (EU1) [25] and European2 (EU2) [26] are 4 major CCs of *M. bovis* that are well described in the literature.

In the last 10 years, it is known that Morocco imported cattle from France, Germany, Spain and, previously, from Canada (Personal communication from 3 different veterinarians working at the national Moroccan Veterinary Services). In addition, there is human immigration from Morocco to Europe and North America. In the present study, the selection process of the countries to include in the second phylogenetic analysis was performed based on cattle (France, Spain, and Germany to Morocco) and human movement (Morocco to Europe and North America), in addition to the preliminary phylogenetic analysis.

The present study aims to determine the genetic relatedness and epidemiological links of *M. bovis* isolates from slaughtered cattle in Morocco using WGS for the first time. Moreover, it also aims to identify *M. bovis* clonal complexes in Morocco and to infer phylogenetic relationships between *M. bovis* isolates from Morocco and from neighboring countries with economic and political links. Investigating *M. bovis* genetic diversity using WGS will provide valuable information that can be used to shape *M. bovis* control strategies in cattle and humans in Morocco. In this study, we performed a preliminary analysis with a representative sample from all the publicly available *M. bovis* isolates, in addition to *M. bovis* isolates from collaborators in Portugal. The objective of the study was to identify genetic links between *M. bovis* cattle isolates and human isolates from Morocco and other countries.

## Material and methods

### Ethical clearance

Gross visible lesions from slaughtered cattle were collected during routine meat inspection at slaughterhouses, therefore no ethical clearance was necessary.

### Sample collection

Gross visible lesions from slaughtered cattle were collected from three abattoirs located in Rabat, El Jadida (western Morocco) and Oujda (Eastern Morocco) during June 2014, from September to December 2014, and from February to July 2015, respectively (Figure 1). The samples included lesions and granulomas found in the animals’ tissues and associated lymph nodes.

**Figure 1.**
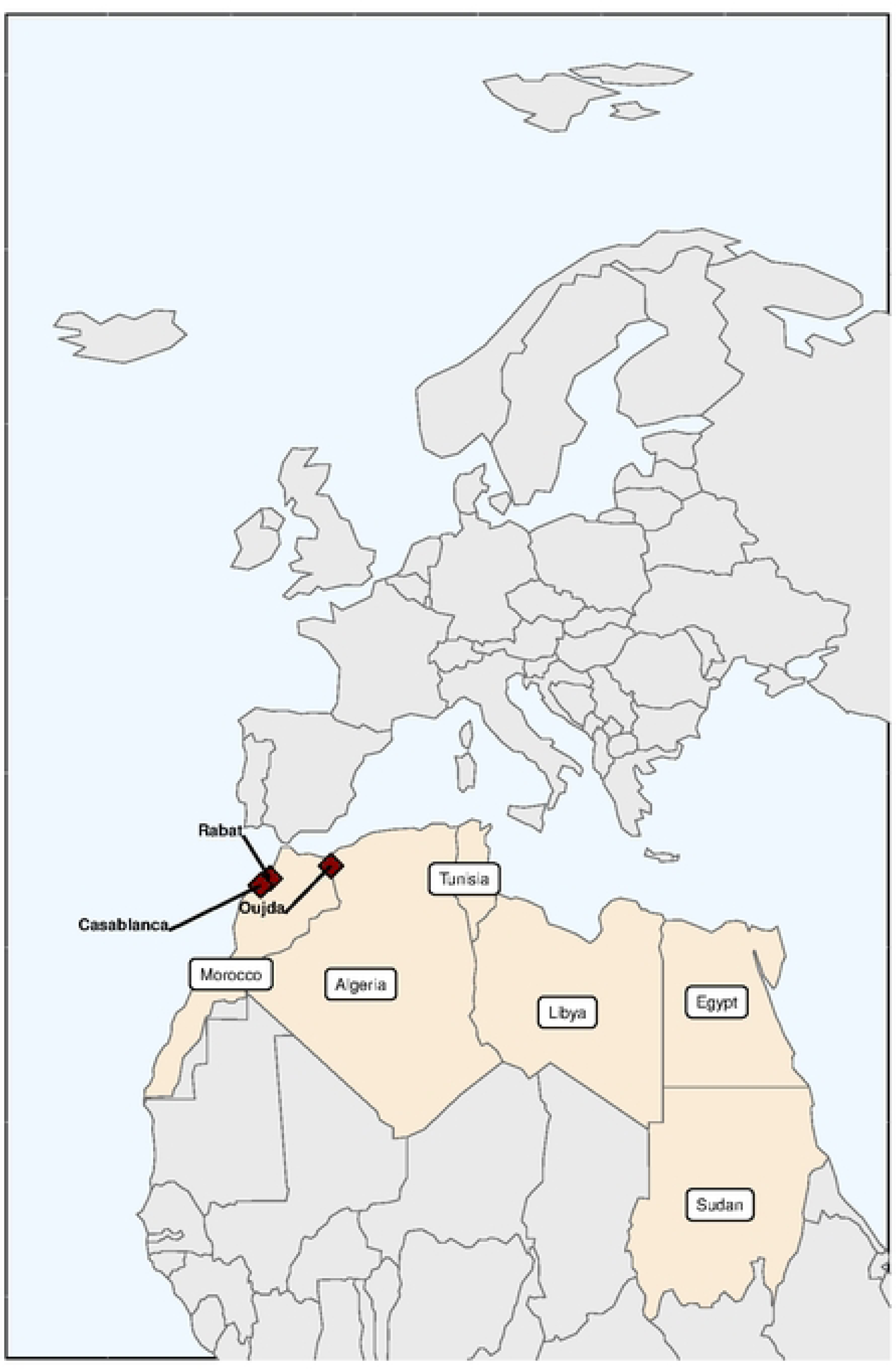
Map showing the slaughterhouse locations from where tissue samples with evidence of bTB-like lesions were collected from cattle. The animals slaughtered at the abattoirs were mostly young bulls and older cows (Table 1). Our samples were composed of more male animals than female (n = 34 vs n = 18). Most of the animals were crossbred (n=40), while a few were labeled as imported (n = 12) (Table 1). Characteristics were not available for 4 animals.

**Table 1:**
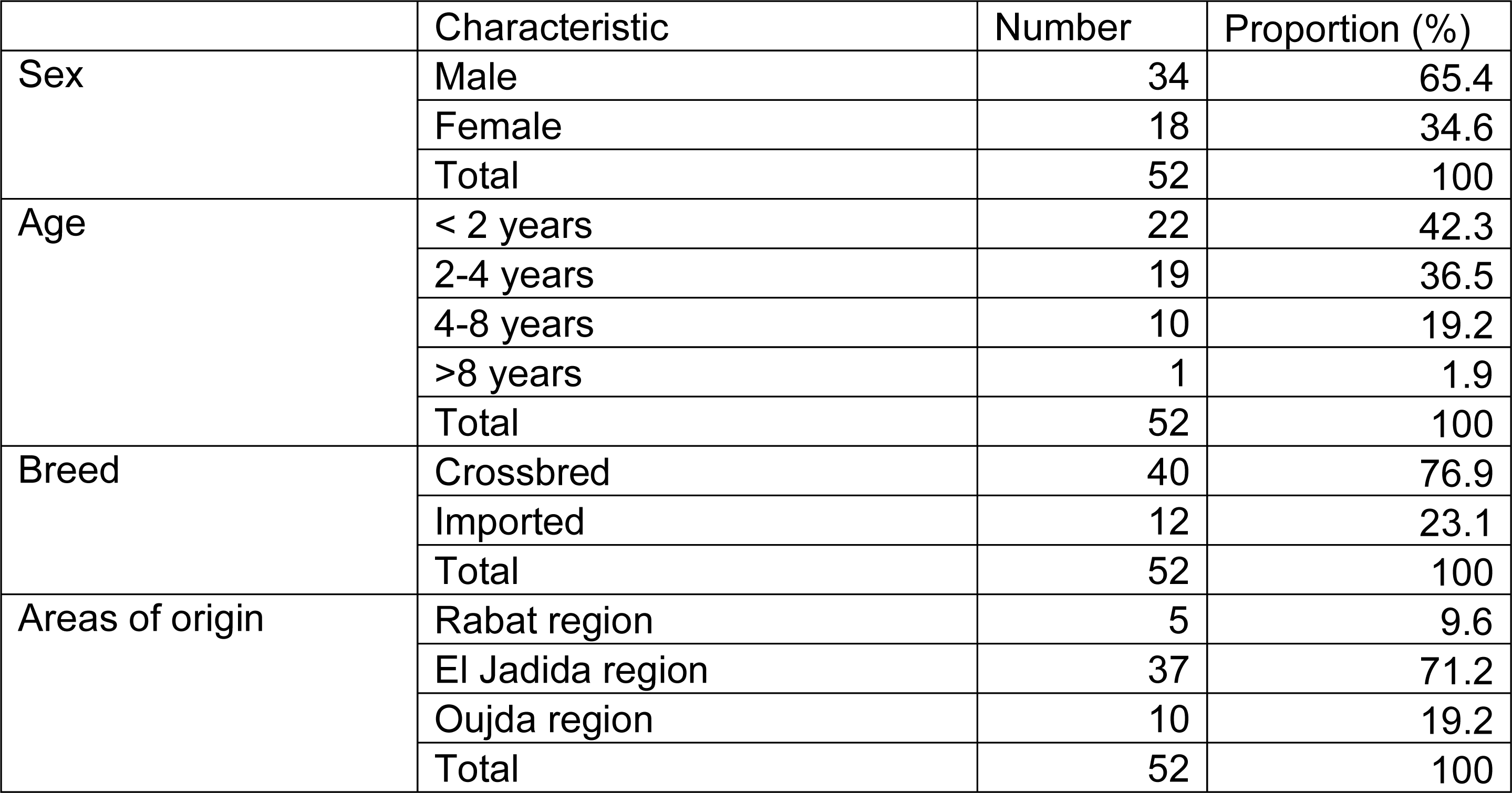
Characteristics of the animals sampled from three slaughterhouses located in El Jadida, Rabat and Oujda during June 2014, from September to December 2014, and from February to July 2015, respectively.

|  | Characteristic | Number | Proportion (%) |
| --- | --- | --- | --- |
| Sex | Male | 34 | 65.4 |
|  | Female | 18 | 34.6 |
|  | Total | 52 | 100 |
| Age | < 2 years | 22 | 42.3 |
|  | 2-4 years | 19 | 36.5 |
|  | 4-8 years | 10 | 19.2 |
|  | >8 years | 1 | 1.9 |
|  | Total | 52 | 100 |
| Breed | Crossbred | 40 | 76.9 |
|  | Imported | 12 | 23.1 |
|  | Total | 52 | 100 |
| Areas of origin | Rabat region | 5 | 9.6 |
|  | El Jadida region | 37 | 71.2 |
|  | Oujda region | 10 | 19.2 |
|  | Total | 52 | 100 |

### Mycobacterial culture

Cattle with gross visible lesions suggesting bTB were sampled, and the lymph nodes and/or other tissues exhibiting lesions from each animal were collected, pooled and stored at -20°C until cultured. Tissue samples were processed in a biosafety level 3 laboratory using sterile surgical instruments.

Frozen samples were thawed overnight at 4°C. Subsequently, 5 g of lesion tissue and adjacent material was mixed with sterilized sand and 10 ml of red phenol. Aliquots of resulting solution (7.5 g) were placed in 15 ml conic tubes with 5 ml of 1N NaOH and incubated at room temperature for 10 minutes. HCl 6N was added for sample neutralization, and the tubes were centrifuged for 25 minutes at 3,000 X g.

The supernatants were discarded, and pellets were distributed in the two pre-tested culture media: Lowenstein Jensen with pyruvate (LJP), and Herrold egg yolk medium. Cultures were incubated for 12 weeks at 37°C. Colonies were sub-cultured on 4 slants of LJP and incubated as previously described for the primary cultures. The cultures were examined weekly for assessment of growth.

### DNA purification

After 4 weeks of incubation at 37°C, the LJP slant cultures were heat inactivated following the USDA heat-killing protocol [27]. Briefly, the colonies from the four LJP slants were pooled, suspended in sterilized distilled water, and heated for 30 minutes at 100°C. After cooling to room temperature, the stocks were stored at -20°C.

The heat-killed mycobacteria were added to the prepared bead beater tubes with 1X Tris-EDTA (TE) buffer and phenol/chloroform/isoamyl alcohol (PCI), the samples were beaten at full speed for 2 minutes using a bead-beater machine (homogenizer). The samples were then spun at 16,000 X g for 5 minutes. The DNA contained in the top aqueous layer was removed (approximately 300 μl) and was added to a 1.5 ml microcentrifuge tube containing 30 μl of a 3M sodium acetate buffer solution. A total of 700 μl of 100% ice-cold ethanol was added to the samples. Samples were cooled at -80°C for 10-15 minutes, then spun at 4°C, 16,000 X g for 15 minutes. The ethanol was discarded and 1 ml ice-cold 70% ethanol was added. The tube was spun at 8,000 X g for 15 seconds and the remaining ethanol was removed. The samples were dried at room temperature and 300 μl of 1X TE buffer was added to resuspend the DNA, which was stored at 4°C for short term storage or -80°C for long term storage [28]. The DNA concentrations were measured using Nanodrop and Qubit 2.0 fluorometer according to manufacturer instructions and subsequently diluted to a starting concentration of 10 ng/μl.

### Whole genome sequencing

A minimum of 20 μL of DNA sample with a minimum concentration of 5 ng/μL was required for sequencing. Libraries were prepared using the Nextera XT Kit (Illumina, Inc., San Diego, CA, USA), and sequencing was performed on an Illumina MiSeq device using 250 bp paired end read chemistry, according to manufacturer’s instructions. Multiple isolates were indexed per lane, providing approximately 50–100X coverage per genome.

### Spoligotyping

Spoligotype profiles were identified *in silico* using WGS data available through the NVSL in-house pipeline vSNP (https://github.com/USDA-VS/vSNP) with the “spoligo” function. The presence or absence of the spacer units in the mycobacterial genome are used to obtain the binary code, which is then cross-referenced against the *Mycobacterium bovis* Spoligotype Database (www.mbovis.org) to obtain the SB codes for each isolate.

### Phylogenetic Analysis

We performed a preliminary analysis of the 55 Moroccan isolates, 125 M. bovis isolates from collaborators in Spain in addition to a representative sample (600 isolates) from all the publicly available *M. bovis* isolates from 34 countries (Supplementary File 1) downloaded from the NCBI Sequence Read Archive.

Raw FASTQ files were analyzed with the National Veterinary Services Laboratory (NVSL) vSNP pipeline [29]. Briefly, the alignment against *M. bovis* AF2122/97 (NC_002945.4) was performed using Burrows-Wheeler Aligner [30] (BWA), and SNPs were called using Freebayes [31]. 80x dept of coverage was targeted. *Mycobacterium caprae* was used as the outgroup. Sites that fell within proline-glutamate (PE) and proline-proline-glutamate (PPE)-polymorphic CG-repetitive sequences (PGRS) were filtered and excluded, as well as SNP positions with a phred-scaled quality (QUAL) score for the alternate nonreference allele lower than 150 or when all positions in a data set had an allele count (AC) equal to 1 when analyzed as a diploid. Integrated Genomics Viewer [32] (IGV) was used to visually validate SNPs, and SNPs with mapping issues or alignment problems were manually filtered. A phylogenetic tree was constructed with RAxML [33] using a GTR-CAT model of substitution and a maximum-likelihood algorithm with the aligned whole-genome SNP sequences. The output from the vSNP pipeline included: an alignment/fasta file containing the concatenated SNP sequences, an Excel table that includes the SNP position (with respect to the reference genome NC_002945.4), annotation and average mapping score of all the SNPs (which are grouped and sorted according to relatedness), and a phylogenetic tree. The accuracy of the phylogenetic tree was confirmed using the validated SNP table. Tree visualization, annotation, and editing were performed with FigTree [34] and iTOL [35]. Clonal complex classification was based on previously published data [36], and on the most recent lineage classification [37].

A total of 780 *M. bovis* genomes were analyzed in the present study, including 56 from Morocco, the rest of the sequences were publicly available and downloaded from NCBI. Thirty-six countries were the focus of this study, including Morocco (Supplementary File 1). The majority of the hosts associated with the analyzed genomes corresponded to cattle (n = 589), human (n=107), and wildlife (n=76) (Supplementary file 1).

## Results

### Spoligotypes of *M. bovis* from Morocco

Twenty-two spoligotyping patterns generated *in sillico* were identified for the Moroccan isolates (Supplementary File 2), of which two were new andwere submitted to the https://www.mbovis.org/ database and assigned new SB numbers (SB2545 and SB2785). The most frequent patterns were SB0121 (n=13, 23.2%), SB0265 (n=12, 21.4%), and SB0120 (n=8, 14.3%).

### SNP map of the Moroccan isolates

A SNP map for the Moroccan isolates was generated. Non-informative SNPs, ambiguous bases, unreliable bases, and gaps were filtered out. The 56 isolates showed a total of 4,410 SNPs compared with the reference strain as shown in Figure 2. Each inner circle represents an isolate, while the outer circle represents the reference strain. The different base substitutions are color coded (green for A, red for T, blue for C and black for G).

**Figure 2.**
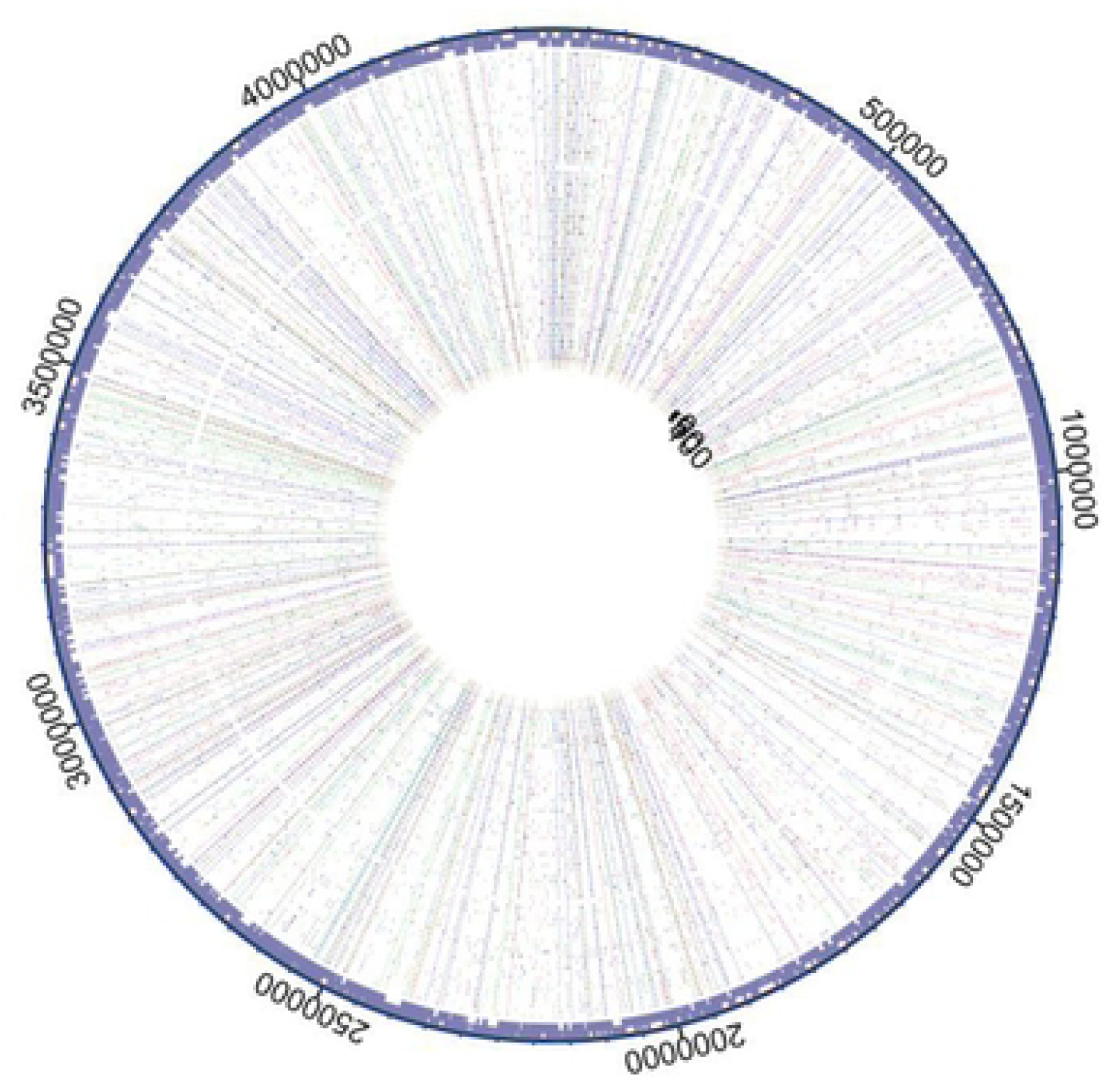
SNP map of the 56 Moroccan M. bovis isolates collected during 2014 and 2015. Each inner circle represents an isolate and the outer circle refers to the reference strain.

### SNP phylogeny of *M. bovis* from Morocco

A total of 784 isolates were analyzed using vSNP, the results identified several genetic groups. Figure 3 shows the overall SNP-based phylogenetic tree including all the isolates analyzed. The countries are color coded, and only 7 countries are emphasized in the key; in addition to Morocco, we included Spain, France, Germany, and Canada, countries where Morocco imported cattle in the last 10 years. Portugal and Algeria were included due to their geographical proximity to Morocco.

**Figure 3.**
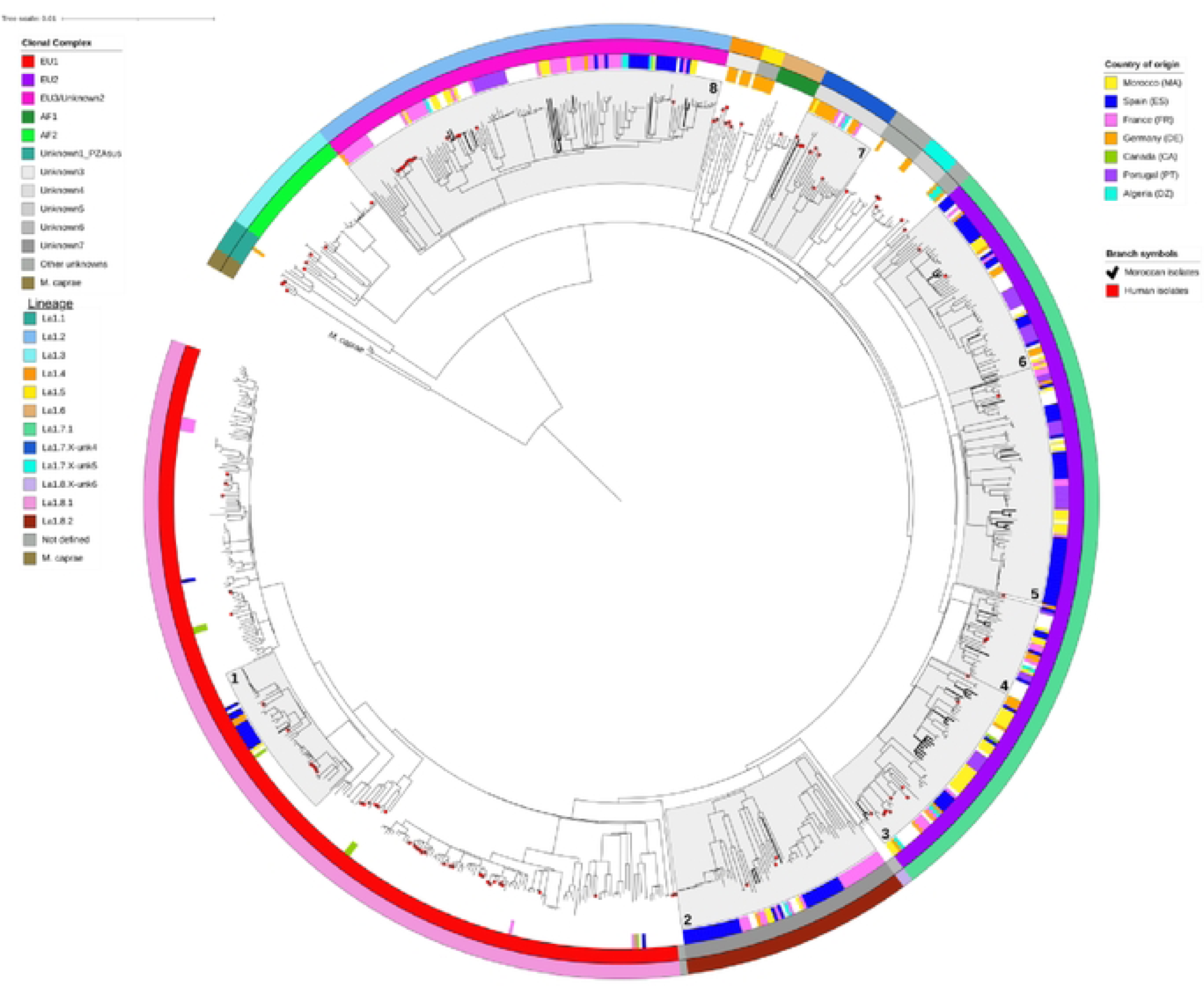
Maximum-likelihood phylogenetic tree with GTR-CAT substitution model presenting all the analyzed *M. bovis* isolates (n=784), with countries (inner circle), clonal complex (middle circle) and lineages (outer circle) color coded. Moroccan isolates are indicated with bold marked tree branches, and human isolates are indicated with a red dot. The areas filled with a light grey shade indicate the groups that included Moroccan isolates.

Most isolates were part of EU1, EU2, EU3-Unknown2 and AF1 clonal complexes, which are equivalent to La 8.1.8, La 1.7.1, La 1.2 and La 1.3 lineages, respectively, in the new clonal complexes’ classification suggested by Zwyer et al [37].

The Moroccan isolates were part of eight genetic groups, based on NVSL classification [9]. Figure 3 shows the different groups highlighted in light grey and numbered from 1 to 8. Moroccan isolates are indicated with bold tree branches and human *M. bovis* are indicated with red marks (Figure 3).

Figure 4 presents each of the eight genetic groups separately corresponding to groups 1 to 8 from Figure 3. Moroccan *M. bovis* sequences showed a variable genetic relatedness with isolates from the other emphasized countries. Panels 1 and 4 from Figure 4 highlight the genetic relationship between *M. bovis* isolates from Morocco and Spain at different degrees, sharing a most recent common ancestor (MRCA) with ≥30 SNPs. In addition, *M. bovis* isolates from Morocco and France also showed a tendency to cluster together, sharing a MRCA with ≥50 SNP (Panels 5, 6 and 8 of Figure 4). Only 7 *M. bovis* from Canada were publicly available at NCBI and included in the analysis, with only 2 falling into the same groups as the Moroccan isolates (Fig 4, panels 1 and 3), however, no close relationships were noted (>100 SNP to a MRCA). Moroccan and Portuguese *M. bovis* isolates showed a close genetic relationship, sharing a MRCA within 50 to 120 SNPs (Fig 4, panels 3,4 and 5). Finally, two *M. bovis* isolates from Morocco shared a MRCA with an isolate from Algeria with 50 SNPs (Fig 4, panel 3).

**Figure 4.**
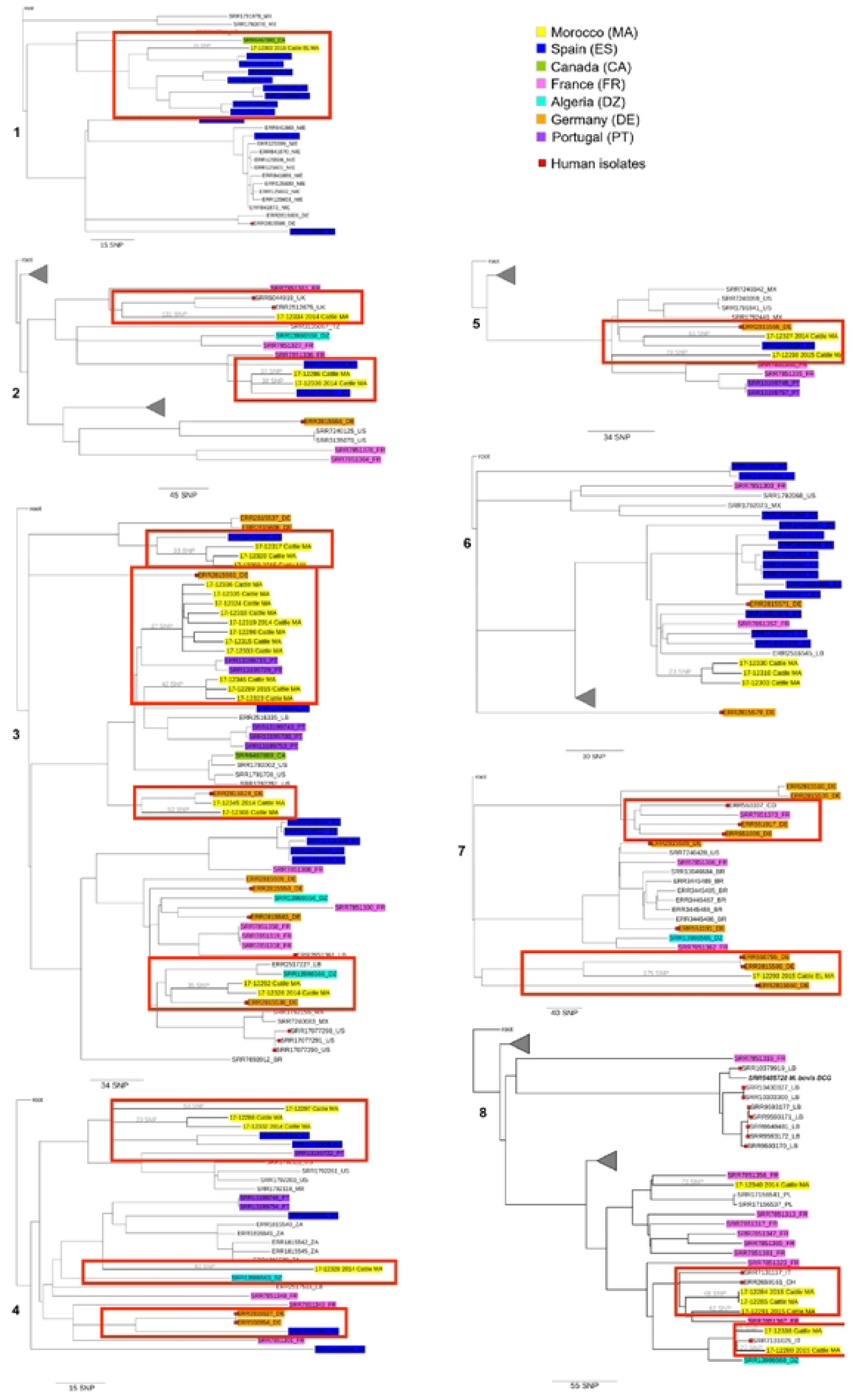
High-resolution maximum-likelihood phylogenetic tree of groups 1-8. The legend indicates country of origin based on highlighted isolate color. The scale bar represents the SNPs branch length. Within this figure, eight panels illustrate genetic relationships among *M. bovis* isolates. **Panel 1** highlights the genetic connection between Moroccan and Spanish *M. bovis* isolates, alongside one of the seven Canadian *M. bovis* isolates analyzed. **Panel 2** reveals genetic relationships between UK human isolates and Moroccan cattle *M. bovis* in the top red box, while the bottom red box showcases the close genetic proximity between *M. bovis* isolates from Spain and Morocco (cattle). **Panel 3** emphasizes genetic proximity between Spanish and Moroccan isolates in the top red box, with the second box displaying clustering between Moroccan and Portuguese *M. bovis*. The third red box in Panel 3 shows a human *M. bovis* isolate from Germany sharing a Most Recent Common Ancestor (MRCA) with a Moroccan cattle *M. bovis* within 20 single nucleotide polymorphisms (SNPs). The bottom box highlights another human *M. bovis* from Germany closely related to Moroccan cattle *M. bovis*, as well as the relationship between an Algerian cattle *M. bovis* and Moroccan cattle *M. bovis*. **Panel 4** demonstrates the close genetic relationship between *M. bovis* isolates from Morocco, Spain, and Portugal. **Panel 5** displays Spanish and Moroccan cattle *M. bovis* isolates and their proximity to German human *M. bovis*. **Panel 7** shows a close genetic relationship between three human German *M. bovis* isolates and one Moroccan cattle *M. bovis*. **Panel 8** illustrates the genetic clustering of human *M. bovis* isolates from Italy and Switzerland alongside Moroccan cattle *M. bovis* in the top red box. The bottom red box highlights the close genetic relationship between a human *M. bovis* isolate from Italy and a cattle *M. bovis* isolate from Morocco, both of which are somewhat genetically related to a cattle *M. bovis* isolate from Algeria.

**Figure 5.**
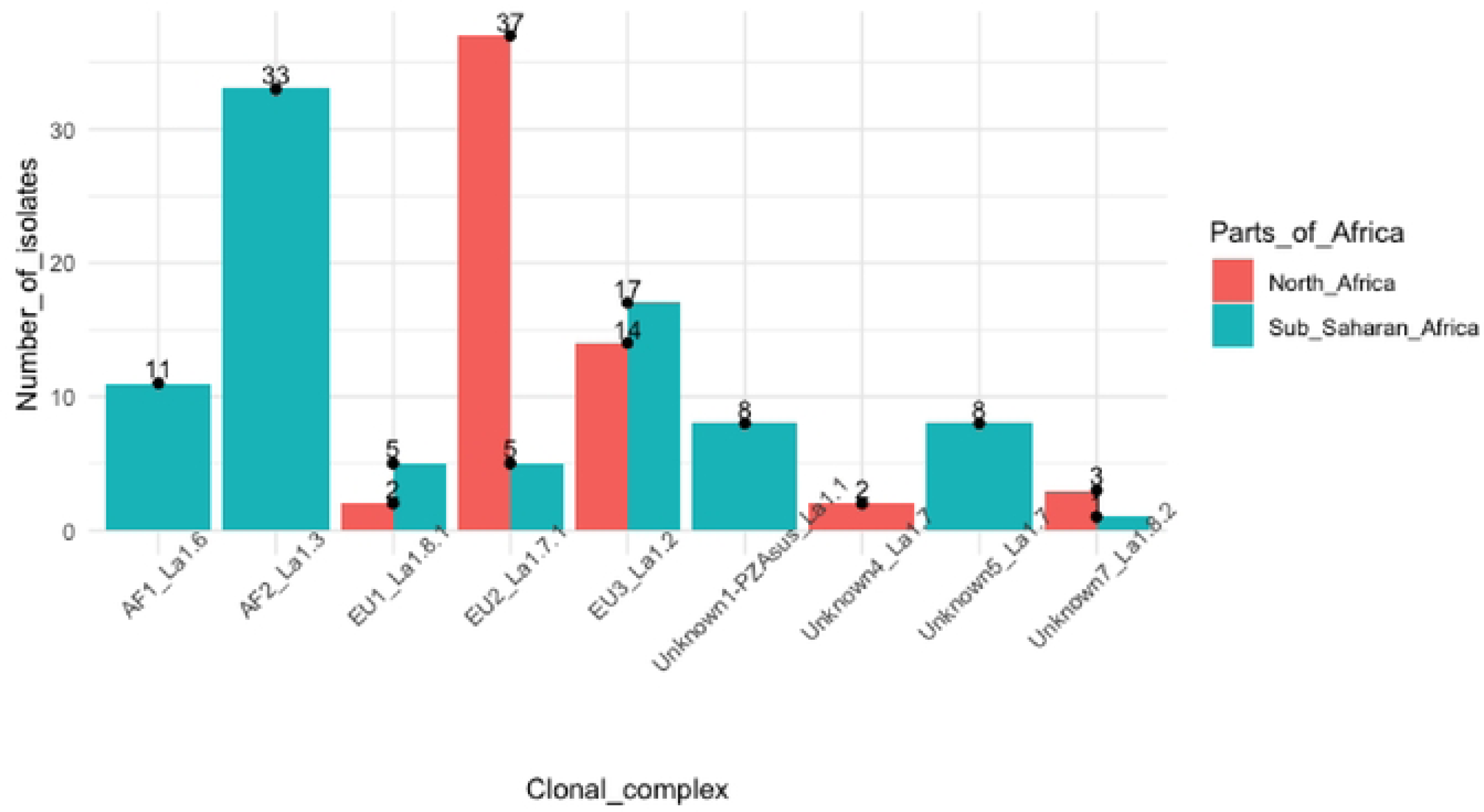
Distribution of Mycobacterium bovis isolates from North Africa and Sub-Saharan Africa based on clonal complexes/lineages. The isolates analyzed here were assessed using the clonal complexes classification. In addition to the most recently described lineage classification, which identified further groups and defined some of the previously unknown groups. *Mycobacterium bovis* sequences from North Africa (Morocco and Algeria) were part of EU1, EU2, EU3, Unknown 1-PZAsus and Unknown 5 clonal complexes, while none of the isolates from North Africa were classified as AF1 and AF2 clonal complexes. All the isolates from AF1 and AF2 clonal complexes were from Sub-Saharan Africa except one (human *M. bovis* isolate collected in Switzerland).

The database analyzed here included 107 *M. bovis* isolates from human patients from Germany, USA, Mexico, UK and Italy. The closest matches of Moroccan cattle to human isolates were from Germany (Fig 4, Panels 3 and 7), the United Kingdom (Fig 4, Panel 2) and Italy (Fig 4, Panel 8), having shared a MRCA with ≥20, ≥130 and ≥16 SNPs, respectively.

## Discussion

The current study represents the first WGS phylogenetic study and SNP analysis of *M. bovis* isolates in Morocco. The analysis included 56 *M. bovis* isolates from three different slaughterhouses in the north of the country and 728 *M. bovis* whole genome sequences downloaded from the NCBI sequence read archive. The results showed that isolates from Morocco are genetically related to isolates from Spain, France, and at a lesser level to isolates from Portugal, Algeria, and Canada. Morocco shares very strong economic and historic ties with Spain, France, and Portugal. Moreover, Morocco has been importing cattle from France, Spain, Canada, and Germany for the last 10 years.

AF1 and AF2 have been shown to be limited to Africa, while EU1 and EU2 have been described in Europe and South America [38]. North Africa has a unique geographical location; it is separated from Sub-Saharan Africa by the Sahara Desert, and it has proximity to Europe, which facilitates commercial trade and human migration. In the current study, *M. bovis* isolates from North Africa were classified as EU1 and EU2 CCs, and none of them fell under AF1 or AF2. This suggests that the Sahara Desert might play a role as a geographical buffer preventing circulation of *M. bovis* between North Africa and Sub-Saharan Africa. This is in line with previous hypothesis based on spoligotype findings in Morocco [15].

The SNP analysis shows *M. bovis* strain similarity between the different countries suggesting that there was a migration of strains perhaps through a combination of human migration (from Morocco to Europe) and cattle movement (from Europe to Morocco). Data from other countries, which are historically and economically linked to Morocco, have not been available. However, we hypothesize that similar links to those observed between Morocco and Spain will exist between Morocco and other European countries, such as Belgium, Germany, Netherlands, Austria and Ireland, from where Morocco has imported cattle for breeding in the past [39]. Furthermore *Mycobacterium bovis* has been identified among wild boar in Morocco [39]; nevertheless, no data are available for the interaction between cattle and wild boar, and the transmission dynamics between the two species. It should be highlighted that wild boar hunting is practiced in Morocco, resulting in human exposure to *M. bovis* and potential transmission from wild boar to humans.

Similarly to what we’re observing in the current study, previous studies in other disease systems have shown phylogenetic similarity between other bacterial pathogens in Morocco and Europe, for example, *Brucella abortus* [40,41]. For instance, *Brucella* strains isolated from three Friesian/Holstein cows in Morocco were found to be *B. abortus* biovar 1, and phylogenomic studies showed that these strains have some similarity with Spanish strains [10].

The analysis included 107 human *M. bovis* isolates from different countries, and genetic similarities were identified between Moroccan cattle *M. bovis* isolates and human *M. bovis* from Italy, Germany, and the UK. During World War I, migration from Morocco to Europe started and it has increased throughout the years, currently there are approximately 5 million Moroccans living abroad, mostly in Europe [43]. This human movement from Morocco to European countries can explain the genetic similarities between cattle *M. bovis* strains from Morocco and human *M. bovis* strains from Europe. Transmission of *M. bovis* from cattle to humans has been documented previously in several countries. A study in Mexico showed that 30.2 % of 533 human tuberculosis patients were infected with *M. bovis* [44]. In Tunisia, two separate studies showed that extrapulmonary TB (EPTB) was caused mainly by *M. bovis* (76% and 77%, respectively) [16,45]. Spoligotyping and MIRU-VNTR analyses of isolates from EPTB patients showed a similarity between the strains isolated from humans and cattle; in addition, most of *M. bovis* TB patients have reported close contact with livestock and consumption of unpasteurized milk and milk products [46]. Zoonotic TB has not been investigated in Morocco, however, bTB has a high individual (18%) and herd (56%) prevalence in cattle [47], and livestock keepers and their families are in very close contact with cattle. These, together with the absence of an official milk pasteurization policy in Morocco, support the likelihood of a high risk of *M. bovis* transmission to humans, particularly in the agriculture sectors of Morocco.

In a low bTB prevalence setting like the USA, the inspection of cattle carcasses at slaughterhouses is followed by sample submission of suspicious bTB cases for postmortem diagnostic tests. Culture and WGS are carried out, and a positive test, results in the condemnation of the carcass. In Morocco, the inspection and identification of bTB-like gross visible lesions is the only element considered when deciding whether to condemn the carcass and the extent of the condemnation. Inspection of cattle carcasses in Morocco are performed by a veterinarian or technician at the slaughterhouse. In addition, information relative to the slaughtered animals is collected, including origin, sex, age, breed, reason for condemnation, the type of condemnation (partial or total) and the organs and parts of the carcass condemned, if the condemnation is partial. The condemned organs and carcasses are then destroyed, and the data are recorded and stored within the veterinary services directory. Even though the inspection method followed is standardized, inspection conditions in some slaughterhouses are not optimal, especially in the rural areas, where access to light at the abattoirs is poor. Such poor conditions can lead to missing small lesions that would have led to the partial condemnation of a carcass under optimal conditions. Inspection of carcasses at the slaughterhouse has proven to be an appropriate surveillance strategy that can prevent contaminated meat and organs from reaching consumers. However, it is not a standalone measure, especially in developing countries, where bTB is endemic, and slaughterhouse conditions are usually poor. In Morocco, most veterinarians and technicians performing the inspection of carcasses do not follow biosecurity procedures, which means that these individuals are directly exposed to the risk of contracting different zoonoses like bTB. In fact, zoonotic TB has been reported among abattoir workers in many countries [48,49]; in Morocco, this issue has not been investigated. Furthermore, even with optimal conditions, sensitivity of slaughterhouse surveillance remains very low (around 44.8%) [50]. Taking into consideration that bTB is endemic in Moroccan cattle, while the zoonotic human health burden is still unknown, the inspection/condemnation method puts slaughterhouse workers, as well as the general human health, at risk of contracting *M. bovis*. In addition, tracking the animals to their herds is not possible because no adequate tracing systems are in place. Having a detailed movement tracking of cattle is an important step in the control of any infectious disease, and it will make a potential test and slaughter intervention to control bTB easier to perform, and more sustainable. Morocco was in the process of launching a campaign to ear tag all the cattle in the country, however, those efforts came to a halt because of the COVID-19 pandemic.

The gold standard for bTB control is test and slaughter, which is based on testing cattle using the simple and comparative tuberculin skin tests followed by the slaughter of test positive animals. Sensitivity to the simple and comparative tuberculin skin tests is not optimal. A low sensitivity will lead to a low detection of infected animals, allowing the persistence of transmitters within the population and leading to a longer time to elimination of the disease. In addition, a mathematical transmission model for bTB in Morocco has shown that the key element regarding bTB elimination is sensitivity of the diagnostic test used [51]. The addition of WGS, when applied to a larger sample size in different areas of Morocco, coupled with strong epidemiological data, can lead to the identification of high transmission areas and potentially sources of transmission in those areas. The application of another screening test in high transmission areas, such as the IFN gamma response assay, which has been shown to have enhanced sensitivity when combined to tuberculin skin test, will help to decrease the false negative results. Lastly, for an optimal bTB control plan, *M. bovis* infection should be investigated in other susceptible hosts within the geographic area, in the case of Morocco, humans, wild boar, small ruminants and camels. Moreover, comparing additional Moroccan isolates with European and African animal and human isolates will provide additional insight regarding the transmission dynamics of *M. bovis* between Europe and Africa, and will aid in preventing future dissemination and re-introductions of the disease.

## Acknowledgments

MVC acknowledges funding from Fundação para a Ciência e a Tecnologia (FCT)/MCTES through national funds (PIDDAC) and co-funding by the European Regional Development Fund (FEDER) of the European Union, through the Lisbon Regional Operational Program and the Competitiveness and Internationalization Operational Program for Portugal 2020 or other programs that may succeed in the scope of project “Colossus: Control Of tubercuLOsiS at the wildlife/livestock interface uSing innovative natUre-based Solutions” (references PTDC/CVT-CVT/29783/2017, LISBOA-01-0145-FEDER-029783, POCI-01-0145-FEDER-029783). Strategic funding from FCT to cE3c and BioISI Research Units (UIDB/00329/2020 and UIDB/04046/2020, respectively) and to the associate laboratory CHANGE (LA/P/0121/2020) are gratefully acknowledged. L.C.M.S acknowledges start-up funds from the University of Georgia Office of Research. We would like to acknowledge Gabrielle Veytsel for creating the map featured in figure 1.

## References

1. Cosivi O, Meslin FX, Daborn CJ, Grange JM. Epidemiology of Mycobacterium bovis infection in animals and humans, with particular reference to Africa. Rev Sci Tech Int Off Epizoot. 1995;14: 733–746.

2. Torgerson PR, Torgerson D. Public health and bovine tuberculosis: what’s all the fuss about? Trends Microbiol. 2010;18: 67–72. doi:info:doi/10.1016/j.tim.2009.11.002

3. Tschopp R, Hattendorf J, Roth F, Choudhoury A, Shaw A, Aseffa A, et al. Cost estimate of bovine tuberculosis to Ethiopia. One Health: The Human-Animal-Environment Interfaces in Emerging Infectious Diseases. Springer; 2012. pp. 249– 268.

4. Schiller I, Waters WR, RayWaters W, Vordermeier HM, Jemmi T, Welsh M, et al. Bovine tuberculosis in Europe from the perspective of an officially tuberculosis free country: trade, surveillance and diagnostics. Vet Microbiol. 2011;151: 153–159. doi:10.1016/j.vetmic.2011.02.039

5. More SJ, Radunz B, Glanville RJ. Lessons learned during the successful eradication of bovine tuberculosis from Australia. Vet Rec. 2015;177: 224–232. doi:10.1136/vr.103163

6. Fitzgerald SD, Kaneene JB. Wildlife Reservoirs of Bovine Tuberculosis Worldwide: Hosts, Pathology, Surveillance, and Control. Vet Pathol. 2013;50: 488–499. doi:10.1177/0300985812467472

7. Garine-Wichatitsky MD, Caron A, Kock R, Tschopp R, Munyeme M, Hofmeyr M, et al. A review of bovine tuberculosis at the wildlife–livestock–human interface in sub-Saharan Africa. Epidemiol Infect. 2013;141: 1342–1356. doi:10.1017/S0950268813000708

8. El Mrini M, Kichou F, Kadiri A, Berrada J, Bouslikhane M, Cordonnier N, et al. Animal tuberculosis due to Mycobacterium bovis in Eurasian wild boar from Morocco. Eur J Wildl Res. 2016;62: 479–482. doi:10.1007/s10344-016-1022-0

9. Orloski K, Robbe-Austerman S, Stuber T, Hench B, Schoenbaum M. Whole Genome Sequencing of Mycobacterium bovis Isolated From Livestock in the United States, 1989–2018. Front Vet Sci. 2018;5. doi:10.3389/fvets.2018.00253

10. Yahyaoui Azami H., Ducrotoy M. J., Bouslikhane M., Hattendorf J., Thrusfield M., Conde-Alvarez R., et al. The prevalence of brucellosis and bovine tuberculosis in ruminants in Sidi Kacem Province, Morocco. PLOS ONE. 2018;In press.

11. Phillips CJC, Foster CRW, Morris PA, Teverson R. The transmission of Mycobacterium bovis infection to cattle. Res Vet Sci. 2003;74: 1–15. doi:10.1016/S0034-5288(02)00145-5

12. Humblet M-F, Boschiroli ML, Saegerman C. Classification of worldwide bovine tuberculosis risk factors in cattle: a stratified approach. Vet Res. 2009;40: 50. doi:10.1051/vetres/2009033

13. Walter WD, Smith R, Vanderklok M, VerCauteren KC. Linking bovine tuberculosis on cattle farms to white-tailed deer and environmental variables using Bayesian hierarchical analysis. PloS One. 2014;9: e90925. doi:10.1371/journal.pone.0090925

14. Ghebremariam MK, Hlokwe T, Rutten VPMG, Allepuz A, Cadmus S, Muwonge A, et al. Genetic profiling of Mycobacterium bovis strains from slaughtered cattle in Eritrea. PLoS Negl Trop Dis. 2018;12: e0006406. doi:10.1371/journal.pntd.0006406

15. Yahyaoui-Azami H, Aboukhassib H, Bouslikhane M, Berrada J, Rami S, Reinhard M, et al. Molecular characterization of bovine tuberculosis strains in two slaughterhouses in Morocco. BMC Vet Res. 2017;13: 272. doi:10.1186/s12917-017-1165-6

16. Siala M, Smaoui S, Taktak W, Hachicha S, Ghorbel A, Marouane C, et al. First-time detection and identification of the Mycobacterium tuberculosis Complex members in extrapulmonary tuberculosis clinical samples in south Tunisia by a single tube tetraplex real-time PCR assay. PLoS Negl Trop Dis. 2017;11: e0005572. doi:10.1371/journal.pntd.0005572

17. Ghavidel M, Mansury D, Nourian K, Ghazvini K. The most common spoligotype of Mycobacterium bovis isolated in the world and the recommended loci for VNTR typing; A systematic review. Microb Pathog. 2018;118: 310–315. doi:10.1016/j.micpath.2018.03.036

18. Guimaraes AMS, Zimpel CK. Mycobacterium bovis: From Genotyping to Genome Sequencing. Microorganisms. 2020;8. doi:10.3390/microorganisms8050667

19. Yassine E, Galiwango R, Ssengooba W, Ashaba F, Joloba ML, Zalwango S, et al. Assessing a transmission network of Mycobacterium tuberculosis in an African city using single nucleotide polymorphism threshold analysis. MicrobiologyOpen. 2021;10: e1211. doi:10.1002/mbo3.1211

20. Glaser L, Carstensen M, Shaw S, Robbe-Austerman S, Wunschmann A, Grear D, et al. Descriptive Epidemiology and Whole Genome Sequencing Analysis for an Outbreak of Bovine Tuberculosis in Beef Cattle and White-Tailed Deer in Northwestern Minnesota. PLoS ONE. 2016;11. doi:10.1371/journal.pone.0145735

21. Trewby H, Wright D, Breadon EL, Lycett SJ, Mallon TR, McCormick C, et al. Use of bacterial whole-genome sequencing to investigate local persistence and spread in bovine tuberculosis. Epidemics. 2016;14: 26–35. doi:10.1016/j.epidem.2015.08.003

22. Sandoval-Azuara SE, Muñiz-Salazar R, Perea-Jacobo R, Robbe-Austerman S, Perera-Ortiz A, López-Valencia G, et al. Whole genome sequencing of Mycobacterium bovis to obtain molecular fingerprints in human and cattle isolates from Baja California, Mexico. Int J Infect Dis. 2017;63: 48–56. doi:10.1016/j.ijid.2017.07.012

23. Müller B, Hilty M, Berg S, Garcia-Pelayo MC, Dale J, Boschiroli ML, et al. African 1, an Epidemiologically Important Clonal Complex of Mycobacterium bovis Dominant in Mali, Nigeria, Cameroon, and Chad. J Bacteriol. 2009;191: 1951–1960. doi:10.1128/JB.01590-08

24. Berg S, Garcia-Pelayo MC, Müller B, Hailu E, Asiimwe B, Kremer K, et al. African 2, a Clonal Complex of Mycobacterium bovis Epidemiologically Important in East Africa. J Bacteriol. 2011;193: 670–678. doi:10.1128/JB.00750-10

25. Smith NH, Berg S, Dale J, Allen A, Rodriguez S, Romero B, et al. European 1: A globally important clonal complex of Mycobacterium bovis. Infect Genet Evol. 2011;11: 1340–1351. doi:10.1016/j.meegid.2011.04.027

26. Rodriguez-Campos S, Schürch AC, Dale J, Lohan AJ, Cunha MV, Botelho A, et al. European 2 – A clonal complex of Mycobacterium bovis dominant in the Iberian Peninsula. Infect Genet Evol. 2012;12: 866–872. doi:10.1016/j.meegid.2011.09.004

27. Ortiz AP, Perea C, Davalos E, Velázquez EF, González KS, Camacho ER, et al. Whole Genome Sequencing Links Mycobacterium bovis From Cattle, Cheese and Humans in Baja California, Mexico. Front Vet Sci. 2021;8: 674307. doi:10.3389/fvets.2021.674307

28. Mycobacteria DNA extraction using bead disruption - Manual method. [cited 13 Dec 2018]. Available: https://www.protocols.io/view/mycobacteria-dna-extraction-using-bead-disruption-nsgdebw

29. USDA. https://github.com/USDA-VS/vSNP.

30. Li H, Durbin R. Fast and accurate short read alignment with Burrows–Wheeler transform. Bioinformatics. 2009;25: 1754–1760. doi:10.1093/bioinformatics/btp324

31. Garrison E, Marth G. Haplotype-based variant detection from short-read sequencing. ArXiv12073907 Q-Bio. 2012 [cited 15 Sep 2021]. Available: http://arxiv.org/abs/1207.3907

32. Robinson JT, Thorvaldsdóttir H, Winckler W, Guttman M, Lander ES, Getz G, et al. Integrative Genomics Viewer. Nat Biotechnol. 2011;29: 24–26. doi:10.1038/nbt.1754

33. Stamatakis A. RAxML version 8: a tool for phylogenetic analysis and post-analysis of large phylogenies | Bioinformatics | Oxford Academic. Available: https://academic.oup.com/bioinformatics/article/30/9/1312/238053

34. Rambaut A. FigTree. Available: http://tree.bio.ed.ac.uk/software/figtree/

35. Letunic I, Bork P. Interactive Tree Of Life (iTOL): an online tool for phylogenetic tree display and annotation. Bioinforma Oxf Engl. 2007;23: 127–128. doi:10.1093/bioinformatics/btl529

36. Loiseau C, Menardo F, Aseffa A, Hailu E, Gumi B, Ameni G, et al. An African origin for Mycobacterium bovis. Evol Med Public Health. 2020;2020: 49–59. doi:10.1093/emph/eoaa005

37. Michaela Zwyer, Çavusoglu C, Giovanni Ghielmetti, Ghielmetti G, Maria Lodovica Pacciarini, et al. A new nomenclature for the livestock-associated Mycobacterium tuberculosis complex based on phylogenomics. [cited 10 Oct 2022]. Available: https://open-research-europe.ec.europa.eu/articles/1-100/v2

38. Ramos DF, Tavares L, Silva PEA da, Dellagostin OA. Molecular typing of Mycobacterium bovis isolates: a review. Braz J Microbiol. 2014;45: 365–372. doi:10.1590/S1517-83822014005000045

39. ONSSA - Bovins. [cited 2 Aug 2018]. Available: http://www.onssa.gov.ma/fr/controle-et-certification-a-limport-export/import/animaux/bovins

40. Taoudi A, Fassi-Fehri MM, Johnson DW, Fagouri S. [Brucella abortus biotypes found in cattle in Morocco: a preliminary study]. Dev Biol Stand. 1983;56: 123–128.

41. Verger Jm, Grayon M. Caractéristiques de 273 souches de Brucella abortus d’origine africaine. Dev Biol Stand. 1984;56: 63–71.

42. El Mrini M, Kichou F, Kadiri A, Berrada J, Bouslikhane M, Cordonnier N, et al. Animal tuberculosis due to Mycobacterium bovis in Eurasian wild boar from Morocco. Eur J Wildl Res. 2016;62: 479–482. doi:10.1007/s10344-016-1022-0

43. Haas H de. Morocco: From Emigration Country to Africa’s Migration Passage to Europe. migrationpolicy.org. 2005 [cited 11 Oct 2022]. Available: https://www.migrationpolicy.org/article/morocco-emigration-country-africas-migration-passage-europe

44. Torres-Gonzalez P, Cervera-Hernandez ME, Martinez-Gamboa A, Garcia-Garcia L, Cruz-Hervert LP, Bobadilla-del Valle M, et al. Human tuberculosis caused by Mycobacterium bovis: a retrospective comparison with Mycobacterium tuberculosis in a Mexican tertiary care centre, 2000–2015. BMC Infect Dis. 2016;16: 657. doi:10.1186/s12879-016-2001-5

45. Ghariani A, Jaouadi T, Smaoui S, Mehiri E, Marouane C, Kammoun S, et al. Diagnosis of lymph node tuberculosis using the GeneXpert MTB/RIF in Tunisia. Int J Mycobacteriology. 2015;4: 270–275. doi:10.1016/j.ijmyco.2015.05.011

46. Siala M, Cassan C, Smaoui S, Kammoun S, Marouane C, Godreuil S, et al. A first insight into genetic diversity of Mycobacterium bovis isolated from extrapulmonary tuberculosis patients in South Tunisia assessed by spoligotyping and MIRU VNTR. PLoS Negl Trop Dis. 2019;13: e0007707. doi:10.1371/journal.pntd.0007707

47. Yahyaoui Azami H, Ducrotoy MJ, Bouslikhane M, Hattendorf J, Thrusfield M, Conde-Álvarez R, et al. The prevalence of brucellosis and bovine tuberculosis in ruminants in Sidi Kacem Province, Morocco. PLoS ONE. 2018;13. doi:10.1371/journal.pone.0203360

48. Robinson P, Morris D, Antic R. Mycobacterium bovis as an occupational hazard in abattoir workers. Aust N Z J Med. 1988;18: 701–703. doi:10.1111/j.1445-5994.1988.tb00156.x

49. Adesokan HK, Jenkins AO, van Soolingen D, Cadmus SIB. Mycobacterium bovis infection in livestock workers in Ibadan, Nigeria: evidence of occupational exposure. Int J Tuberc Lung Dis Off J Int Union Tuberc Lung Dis. 2012;16: 1388– 1392. doi:10.5588/ijtld.12.0109

50. Garcia-Saenz A, Napp S, Lopez S, Casal J, Allepuz A. Estimation of the individual slaughterhouse surveillance sensitivity for bovine tuberculosis in Catalonia (North-Eastern Spain). Prev Vet Med. 2015;121: 332–337. doi:10.1016/j.prevetmed.2015.08.008

51. Abakar MF, Yahyaoui Azami H, Bless PJ, Crump L, Lohmann P, Laager M, et al. Transmission dynamics and elimination potential of zoonotic tuberculosis in morocco. PLoS Negl Trop Dis. 2017;11: e0005214. doi:10.1371/journal.pntd.0005214

